# Torsion of the heart tube by shortage of progenitor cells : identification of *Greb1l* as a genetic determinant of criss-cross hearts in mice

**DOI:** 10.1101/2023.05.11.540418

**Authors:** Ségolène Bernheim, Adrien Borgel, Jean-François Le Garrec, Emeline Perthame, Audrey Desgrange, Wojciech Krezel, Francesca Raimondi, Damien Bonnet, Lucile Houyel, Sigolène M. Meilhac

**Author notes:** Correspondence to Sigolène Meilhac.

## Abstract

Despite their burden and impact, most congenital defects remain poorly understood by lack of knowledge of the embryological mechanisms. Here, we identify *Greb1l* mutants as the first mouse model of criss-cross heart. Based on 3D quantifications of shape changes, we demonstrate that torsion of the atrioventricular canal occurs together with supero-inferior ventricles at E10.5, after heart looping. Mutants phenocopy specific features of partial deficiency in retinoic acid signalling, suggesting that GREB1L is a novel modulator of this signalling. Spatio-temporal gene mapping and cross-correlated transcriptomic analyses further reveal the role of *Greb1l* in maintaining a pool of precursor cells during heart tube elongation, by controlling ribosome biogenesis and cell differentiation. Growth arrest and malposition of the outflow tract are predictive of abnormal tube remodelling in mutants. Our work on a rare cardiac malformation opens novel perspectives on the origin of a broader spectrum of congenital defects associated with *GREB1L* in humans.

**Highlights:**

- *Greb1l* inactivation is the first model of criss-cross heart
- Growth arrest of the outflow tract and reduced pole distance are predictive of the torsion of the atrioventricular canal, and also account for associated defects of supero-inferior ventricles and malposition of the great vessels
- Ventricle position needs to be maintained after heart looping
- GREB1L, which is associated in humans with a spectrum of congenital defects, is required to maintain precursor cells, by promoting ribosome biogenesis and restricting cell differentiation.
- GREB1L is a novel factor involved in retinoic acid signalling.

**In Brief:** *GREB1L* is associated with a spectrum of congenital defects in humans. Bernheim et al now uncover its function in maintaining a reservoir of precursor cells. Inactivation of *Greb1l* in the mouse impairs the elongation of the heart tube leading to criss-cross heart with supero-inferior ventricles.

## Introduction

Tubes are basic structural components of several organs, transporting metabolites. Yet, they acquire specific shapes for specific functions. The heart initially forms as a straight tube in the early embryo, sustaining a single blood flow. Cardiac chambers are added sequentially along the tube axis, so that emergence of the double blood circulation during mammalian development is associated with remodelling of the heart tube. An important step is the looping of the heart tube, which positions cardiac chambers, before the partitioning of the left and right cardiac pumps (Desgrange et al., 2018).

Failure to establish the double blood circulation during development is a life-threatening condition, by impairment of the overall blood oxygenation level. Collectively congenital heart defects are common, affecting 1% of births, but individually may correspond to rare disorders, requiring expert diagnosis. Currently, most structural congenital heart defects have an unknown genetic origin (Houyel and Meilhac, 2021). Criss-cross heart is an example of a very rare congenital heart defect (frequency of 1/125 000 births), with unknown genetic and embryological origins (Sanders, 2003). Criss-cross heart is defined by abnormal atrioventricular connections, such that the mitral and tricuspid valves are not localised on the same plane, resulting in crossing of the left and right blood flows (Anderson et al., 1974). Criss-cross heart is almost always associated with other cardiac defects, including malalignments (supero-inferior ventricles, malposition of the great arteries such as double outlet right ventricle), and right-sided obstruction (pulmonary stenosis, right ventricle hypoplasia) (Anderson, 1982; Freedom et al., 1978; Manuel et al., 2018). Based on anatomical observations, it has been hypothesised (1) that criss-cross heart results from a twist of ventricles following septation (Anderson, 1982; Anderson et al., 1974; Seo et al., 1992) and (2) that ventricle malposition reflects defective heart looping (Jacobs et al., 2007; Van Praagh et al., 1964). Yet, these hypotheses have never been tested experimentally, by lack of a mouse model.

Remodelling of the heart tube has been investigated in animal models. Shape changes are concomitant with the elongation of the tube, by ingression of heart precursors at both poles. In the mouse, elongation of the tube occurs, while the distance between the poles remains fixed. This is the basis of a buckling mechanism, central to the process of heart looping at E8.5, thus highlighting the importance of mechanical constraints (Le Garrec et al., 2017). Cell ingression continues after heart looping, and is important for the elongation of the outflow tract (Francou et al., 2014; Ramsbottom et al., 2014). But how this later ingression affects the shape of the whole heart tube has been overlooked. Cell ingression is coupled with cell differentiation : the heart tube is composed of a myocardium layer, whereas cells in continuity with the tube, in the dorsal pericardial wall, are a reservoir of undifferentiated cardiac precursors, referred to as the second heart field (Kelly et al., 2001). Markers associated with cardiomyocyte differentiation are now well documented by recent profiling of single embryonic cardiac cells (Tyser et al., 2021; Zhang et al.). Based on clonal analyses and genetic tracing, the progressive ingression of cells fated to specific cardiac regions has been reconstructed (Meilhac and Buckingham, 2018) : the posterior second heart field, for example, contributes to both poles of the heart, including the atria and the inferior outflow tract, the source of the pulmonary trunk (Bertrand et al., 2011; Lescroart et al., 2012). Cell ingression and heart tube elongation are regulated by multiple factors, including cardiomyocyte differentiation factors, cell proliferation factors as well as regulators of epithelial architecture (Lyons et al., 1995; Park et al., 2006; Francou et al., 2014; Palmquist-Gomes and Meilhac, 2022). Cell ingression is also modulated by patterning cues, which provide dynamic positional information at specific stages. For example, transient Nodal signalling in the left heart field biases cell behaviour to control the asymmetric remodelling of the heart tube during heart looping (Desgrange et al., 2020). Retinoic acid (RA), a metabolite of vitamin A, patterns the heart field in the antero-posterior axis and controls its size, with a dose-sensitive effect (Bernheim and Meilhac, 2020; De Bono et al., 2018; Duong et al., 2021). Whereas the regulation of the dose of RA has been well studied, involving a few synthesising and degrading enzymes, as well as antagonistic signalling by FGF, how RA signalling is perceived by cells remains incompletely understood. RA has been shown to bind RAR nuclear receptors to activate transcription of target genes via RARE regulatory sequences. However, a recent study has highlighted that only 50% of RA targets follow this canonical mechanism (Berenguer et al., 2020), suggesting the existence of alternative effectors.

*Greb1l* has been identified in embryonic stem (ES) cells as a direct target of RXR receptors, activated upon RA stimulation (Simandi et al., 2016). In humans, *GREB1L* is associated with a broad spectrum of congenital defects, including renal agenesis, hearing impairment, thymic, genital and skeletal anomalies (Schrauwen et al., 2020) : 49 de novo or inherited deleterious dominant alterations of *GREB1L* have been identified, underlining its importance as a disease-causing gene. In the mouse, knock-out of *Greb1l* is homozygous lethal at fetal stages, reproducing several human defects, and with additional defects in the heart and head (De Tomasi et al., 2017). However, pathological mechanisms upon *Greb1l* inactivation have remained enigmatic, by lack of knowledge of the function of the encoded protein, as well as of the cells in which it is active.

Here, we identify *Greb1l* mutants as the first mouse model of criss-cross heart with supero-inferior ventricles. Based on quantitative analyses of gene expression and shape changes, taking into account the spatio-temporal dynamics, we address the embryological origin of criss-cross heart. Our results demonstrate that *Greb1l* is not required for heart looping, but for outflow tract growth thereafter, thus maintaining the overall heart tube shape. With tailored transcriptomic analyses, we further show that *Greb1l* is required to maintain a pool of cardiac precursors, regulating not only cardiomyocyte differentiation, but also ribosome biogenesis. We find that *Greb1l* mutants phenocopy partial deficiency in RA signalling, without affecting RARE target genes. Overall, our work identifies *Greb1l* as a genetic determinant of criss-cross heart in mice and unravel pathological mechanisms, potentially relevant to other congenital defects.

## Results

### *Greb1l* mouse mutants model human criss-cross heart

We analysed in more detail *Greb1l* mouse mutants, which were previously reported to have heart defects with chamber malalignment (De Tomasi et al., 2017). *Greb1l*fetuses were collected at E13.5, before lethality. In 3D images of the heart, we observed an abnormal supero-inferior (i.e. cranio-caudal) position of the ventricles, compared to the left-right position in controls (Fig. 1A-B). The atria were normally positioned and atrioventricular connexions crossed each other (Fig. 1F-G). The aorta and pulmonary trunk were correctly septated but parallel and both connected to the right ventricle (Fig. 1A-B, F-G, Fig. S1N-O), indicative of double outlet right ventricle. Crossing of atrioventricular connexions is a very rare feature of congenital heart defects, diagnostic of criss-cross heart (Fig. 1D-E), not reported in heterotaxy (Lin et al., 2014), which is classically associated with cardiac chamber malalignment. Since criss-cross heart has never been reported in mice, we quantified the heart structure comparatively in *Greb1l* mouse mutants and patients with the same phenotype, i.e. criss-cross heart with supero-inferior ventricles (Fig. 1C). The planes of the left and right atrioventricular valves were found at a 74°±10 angle in *Greb1l* mouse mutants, which is non-significantly different from the 57°±17 angle in diseased patients, and significantly different from the null value in both mouse and patient controls (Fig. 1H). Consistently, the planes of the interatrial and interventricular septa were found at an average 60° angle in *Greb1l* mouse mutants and 68° in diseased patients, whereas they are parallel in control hearts (Fig. 1I). Finally, the 3D orientation of the right ventricle-left ventricle axis was quantified in the endogenous context of the thoracic cavity. In both *Greb1l* mouse mutants and diseased patients, the ventricles clearly aligned with the cranio-caudal axis, as distinct from control hearts (Fig. 1J). We have thus identified the first mouse model of the criss-cross heart disease, with a 100% penetrance, opening the possibility to investigate the etiology of this poorly understood congenital heart defect.

**Figure 1.**
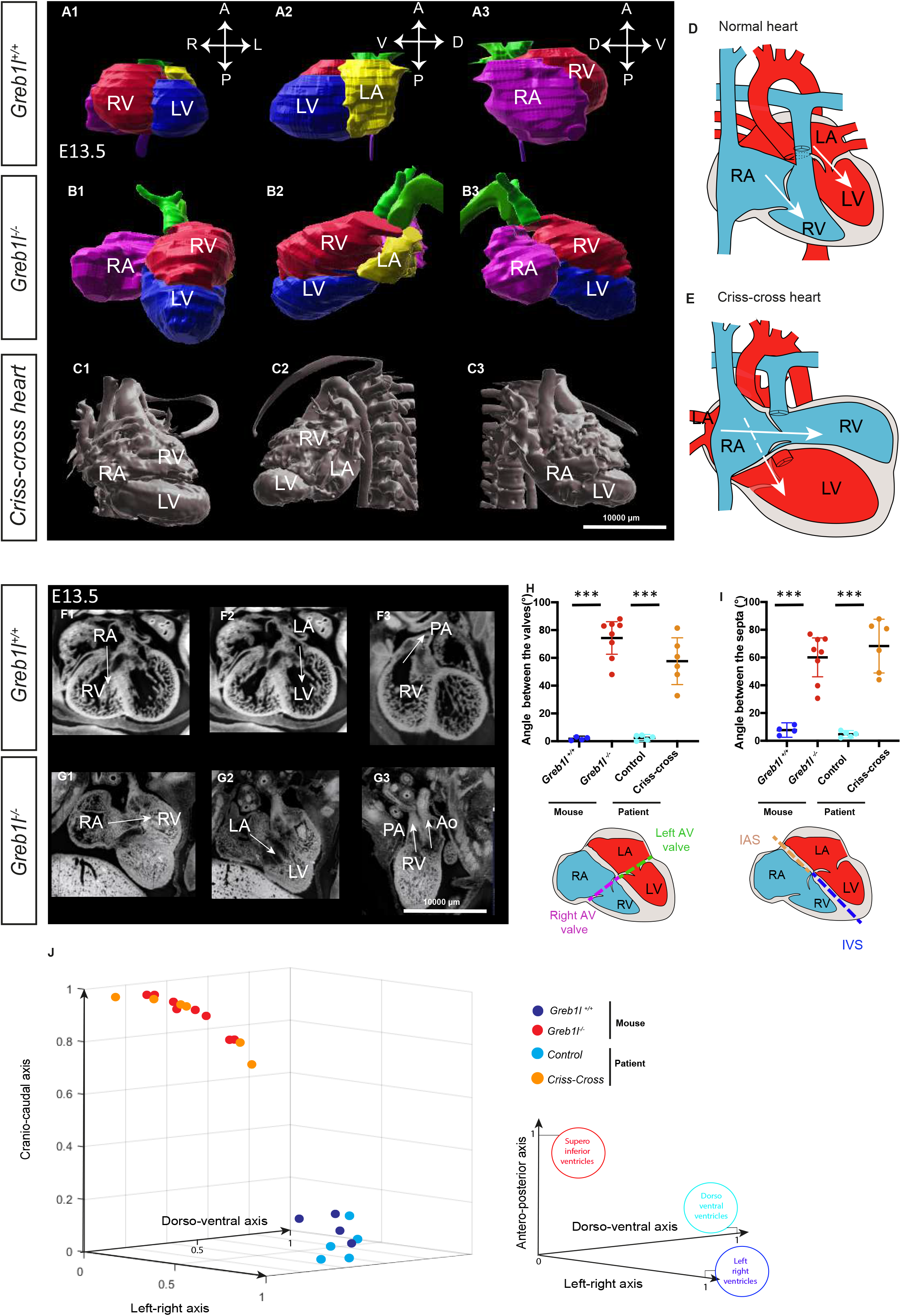
**Quantification of structural heart defects in *Greb1l* mouse mutants at E13.5 by comparison with human criss-cross heart.** (A-B) 3D segmented heart from HREM in controls (A) and *Greb1l^-/-^* mutants (B) in ventral (A1-B1), left-(A2-B2) and right-side (A3-B3) views. (C) 3D segmented heart from a patient CT-scan with criss-cross heart and supero-inferior ventricles. (D-E) Schema of the structure of criss-cross heart (E) compared to normal (D). (F-G) Coronal sections of control (F) and *Greb1l* mutant (G) hearts, showing atrioventricular (arrows, F1-G2) and arterio-ventricular (arrows, F3-G3) connexions. (H) Quantification of the angle between the left and right atrioventricular (AV) valves in mouse and patient hearts. (I) Quantification of the angle between the interatrial (IAS) and interventricular (IVS) septa in mouse and patient hearts. ***p-value<0.001 (One-way ANOVA, n=4 *Greb1l^+/+^*, 8 *Greb1l^-/-^*, 5 control patients, 6 diseased patients). (J) 3D coordinates of the RV/LV axis in mice and patients displaying criss-cross hearts with supero-inferior ventricles compared to controls. Means and standard deviations are shown. A, anterior; Ao, aorta; D, dorsal; L, left; LA, left atrium; LV, left ventricle; n, number of observations; P, posterior; PA, pulmonary artery; R, right; RA, right atrium; RV, right ventricle; V, ventral. See also Video S1.

### *Greb1l* is expressed in cardiac precursors

Despite the involvement of *GREB1L* in several congenital defects (Schrauwen et al., 2020), its expression profile during development has been poorly characterized. We imaged this in 3D at sequential stages of heart development. In addition to expression in the neural tube, endoderm and posterior paraxial mesoderm, we detected *Greb1l* in the lateral plate mesoderm, including cardiac cells, from the late headfold stage (Fig. 2A-B and not shown). Cardiac expression is transient, maximal at E8.5c-d, and by E9.5 it was no longer discernible (Fig. 2B-D). Within the forming heart tube, *Greb1l* was expressed in the myocardium and endocardium (Fig. 2B4-C4). However, expression was found 7-fold higher in the heart field compared to the heart tube (Fig. 2E). These observations are supported by published single cell transcriptomics of cardiac cells (Tyser et al., 2021), where *Greb1l* levels are higher in myocardial precursor clusters Me5 (juxta-cardiac field) and Me7 (second heart field) and higher at crescent stages. *Greb1l* positive cells show increased proliferation capacity compared to *Greb1l* negative cells, with a lower proportion of cells in G1 (Fig. 2F). We identified in the single cell dataset the genes which significantly correlated with variations in *Greb1l* expression levels. The top genes most significantly negatively correlated with *Greb1l* correspond to the cardiomyocyte differentiation genes *Actc1*, *Myl4/7*, *Tmp1*, *Tnnt2* and *Tnni1* (Fig. 2G). Overall, we conclude that *Greb1l* is expressed in early cardiac precursors, before and at the time of heart tube formation.

**Figure 2.**
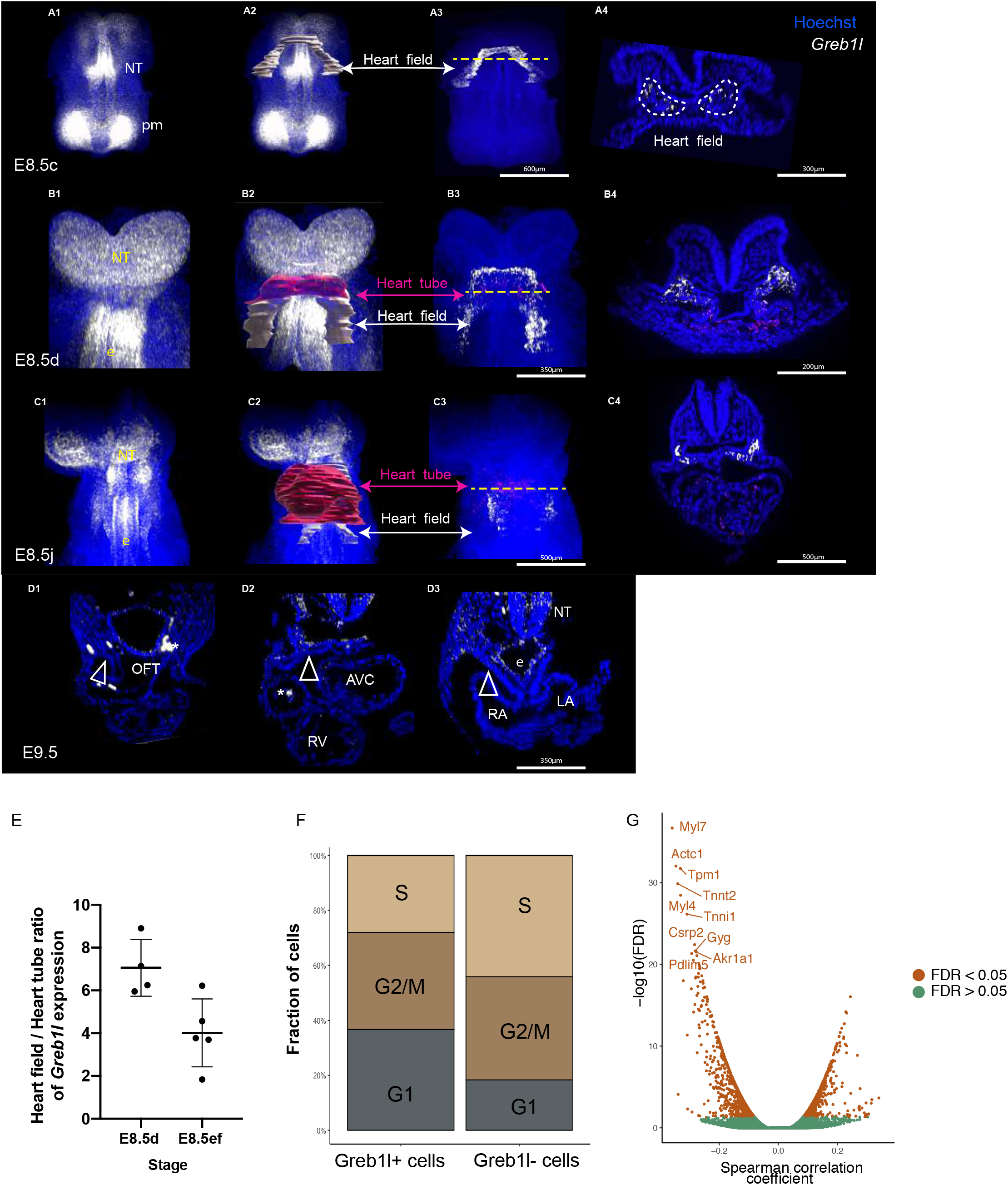
**Expression pattern of *Greb1l* at E8.5.** (A-C) Expression of *Greb1l* (white) detected by whole mount RNAscope ISH in control embryos at E8.5c (A) E8.5d (B), E8.5j (C) and shown in a ventral view (A1-C3) and in transversal sections (A4-C4). Segmentation of the cardiac tissue is indicated in A2-C2, distinguishing the heart field (white) and heart tube (red). Expression of *Greb1l* within the segmented domains is shown in A3-C3, with the same colour code. (D) Expression of *Greb1l* at E9.5 in serial transverse sections, from anterior to posterior (n=3). Arrows point to the negative heart field. Asterisks, unspecific aggregates. (E) Corresponding quantification of the relative expression of *Greb1l* in the heart field and heart tube at the indicated stages (n=4 E8.5d, 5 E8.5e-f). (F) Fraction of wild-type embryonic cardiac cells (Tyser et al., 2021 dataset) assigned transcriptionally to the different cell cycle phases (n=1383 and 485 *Greb1l*+ and *Greb1l*-cells, respectively). (G) Volcano plot for co-expression analysis with *Greb1l* expression levels. Spearman correlation between genes of the dataset (Tyser et al., 2021) and *Greb1l* expression levels in single embryonic cardiac cells is represented against its significance (n=1868 cells, Benjamini-Hochberg corrected p-value < 0.05 in brown and > 0.05 in green). AVC, atrioventricular canal; e, endoderm; FDR, False Discovery Rate; LA, left atrium; LV, left ventricle; NT, neural tube; OFT, outflow tract; pm, paraxial mesoderm; RA, right atrium ; RV, right ventricle.

### *Greb1l* is not required for heart looping

We noticed similarities between *Greb1l* and *Nodal* (Desgrange et al., 2020): both genes are expressed in cardiomyocyte precursors before the formation of the heart tube and both are required for ventricle position. We thus hypothesised that *Greb1l* could be required, similar to *Nodal*, for heart looping, the process during which the right ventricle, initially formed cranially, acquires its position on the right side of the left ventricle. We quantified in *Greb1l* mutants the 3D shape of the heart tube at E9.5, when heart looping is complete. We observed a right-sided position of the right ventricle at E9.5 (Fig. 3A-B), not significantly different from controls (Fig. 3G), as well as normal patterning of the heart tube, as seen with the cardiac chamber marker *Nppa* (Fig. 3C-D) and left markers *Wnt11* and *Bmp2* (Fig. 3E-F). We have shown previously that heart looping results from a buckling mechanism (Le Garrec et al., 2017) : the associated parameters of heart tube length and distance between the poles did not significantly change in *Greb1l* mutants at E9.5 (Fig. 3 H-J). In an independent mutant line that targets a different exon, *Greb1l^ex17/ex17^*, we again did not detect anomalies of heart looping (Fig. S1A, D). Taken together, our 3D analyses show that *Greb1l* is not required for heart looping.

**Figure 3.**
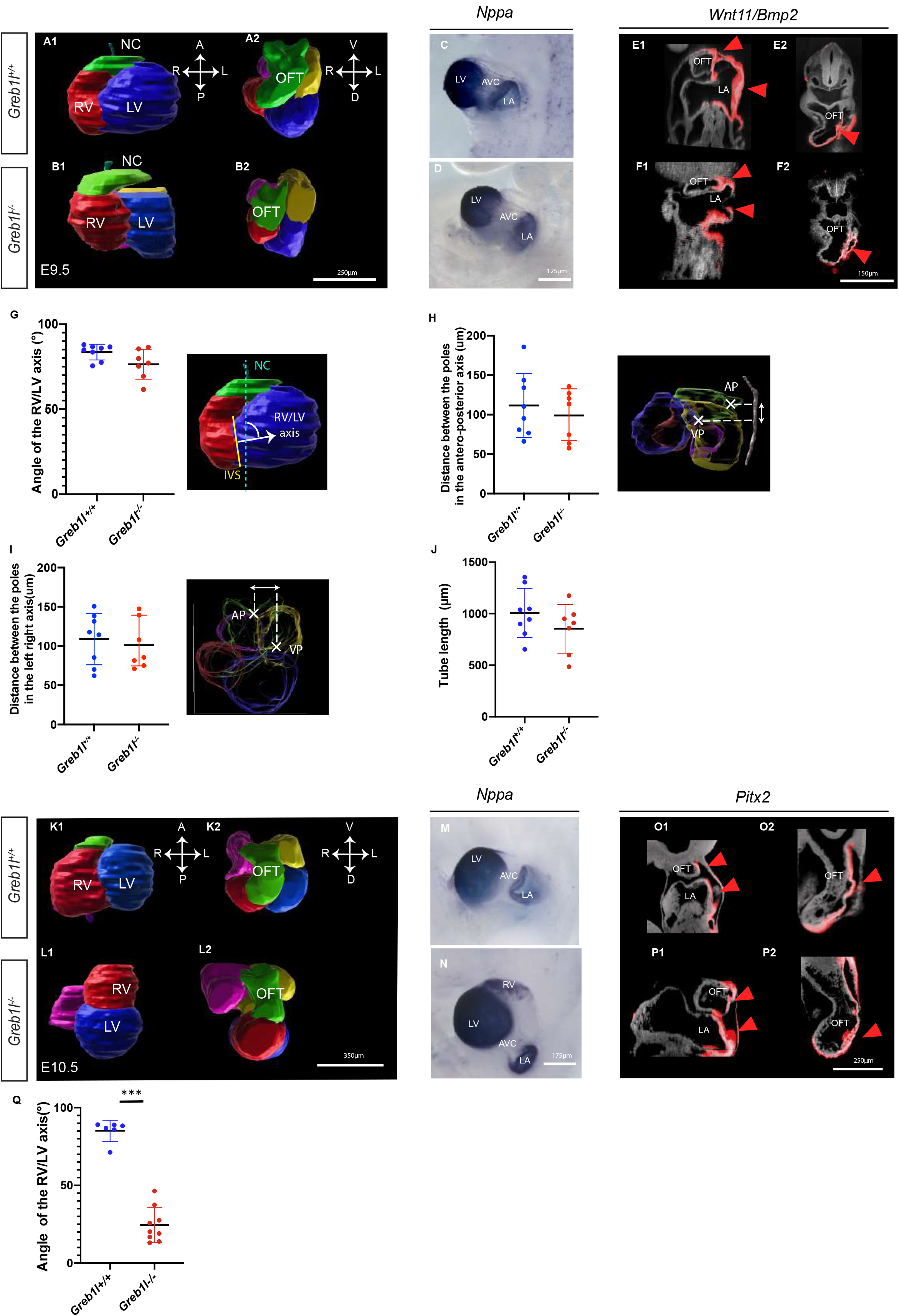
**Geometry and patterning of the heart tube in *Greb1l* mutant embryos at E9.5-E10.5.** (A-B) 3D segmented heart at E9.5 in controls (A) and *Greb1l^-/-^* mutants (B), shown in frontal (A1-B1) and cranial (A2-B2) views. The notochord (NC) is used as the midline reference axis. (C-D) In situ hybridization of *Nppa*, labelling cardiac chambers, in controls (C, n=4) and *Greb1l^ex17/ex17^* mutants (D, n=4) at E9.5, in a left-sided view. (E-F) HREM coronal (E1-F1) and transverse (E2-F2) sections of controls (E, n=4) and *Greb1l^-/-^* mutants (F, n=4). The left-sided expression of *Wnt11* and *Bmp2* is in red, the histology in grey. (G-J) Corresponding quantifications of the RV/LV axis (G), antero-posterior (H) and left-right (I) distance between the heart tube poles, and tube length (J) in controls (n=8) and *Greb1l^-/-^* mutants (n=7) at E9.5 (non-significantly different t-test). (K-L) 3D segmented heart at E10.5 in controls (K) and *Greb1l^-/-^* mutants (L), shown in frontal (K1-L1) and cranial (K2-L2) views. (M-N) In situ hybridization of *Nppa* in controls (M, n=3) and *Greb1l^-/-^* mutants (N, n=5) at E10.5. (O-P) HREM coronal (O1-P1) and transverse (O2-P2) sections of controls (O1, n=3) and mutants (P, n=4). The left-sided expression of *Pitx2* is in red. (Q) Corresponding quantification of the RV/LV axis in controls (n=8) and *Greb1l^-/-^* mutants (n= 9) at E10.5. ***p<0.001 (Mann-Whitney test). Means and standard deviations are shown. A, anterior; AP, arterial pole; AVC, atrioventricular canal; D, dorsal; IVS, interventricular septum; L, left; LA, left atrium; LV, left ventricle; NC, notochord; R, right; RV, right ventricle; OFT, outflow tract; P, posterior; V, ventral; VP, venous pole. See also Figure S1, Videos S2-S3.

### *Greb1l* is required to maintain ventricle position after heart looping

Given the correct position of ventricles in the embryonic E9.5 heart tube of *Greb1l* mutants, we analysed the kinetics of heart shape changes until the criss-cross heart phenotype seen at E13.5. We observed a dramatic change at E10.5, with no further deterioration at subsequent stages (Fig. S1F-O). This kinetics was similarly observed in the two mutant lines, with 100% penetrance (Fig. S1A-E). At E10.5, the ventricles of *Greb1l* mutants have become supero-inferior (Fig. 3K-L, Q), whereas patterning of the heart, with the cardiac chamber marker *Nppa* (Fig. 3M-N) and left marker *Pitx2* (Fig. 3 O-P) was unaffected. Thus, *Greb1l* is required to maintain ventricle position after heart looping, without affecting heart tube patterning.

### Growth arrest of the outflow tract and reduction of pole distance are predictive of ventricle position

Changes in ventricle position in *Greb1l* mutants was accompanied by additional defects at E10.5. The distance between the heart tube poles was significantly reduced in *Greb1l* mutants, in both the left-right and cranio-caudal axes (Fig. 4A-B). The heart tube length was significantly decreased, as a result of a significant shortening of the outflow tract (Fig. 4C-D), whereas the rest of the tube had a constant length (888±175µm in 6 controls compared to 841±95µm in 8 mutants, p=0.5, t-test). Cushions of the atrioventricular canal, which are parallel and supero-inferior in control hearts, had a spiralling pattern in *Greb1l* mutants (Fig. 4E-F). Spiralling of cushions is normally a feature of the outflow tract (Kramer, 1942), reflecting its rightward rotation (Bajolle et al., 2006). We detect here for the first time a case of spiralling of atrioventricular cushions, which thus indicates abnormal leftward rotation of the atrioventricular canal in *Greb1l* mutants. Given that cushions are precursors of the valves, rotation of atrioventricular cushions is consistent with the later phenotype of criss-cross heart. Finally, outflow cushions, had spiralling defects in *Greb1l* mutants (absence of spiralling or abnormal direction of rotation, n=5) (Fig. S1P-U). Outflow tract defects (shortening and abnormal spiralling) are consistent with the later phenotype of double outlet right ventricle.

**Figure 4.**
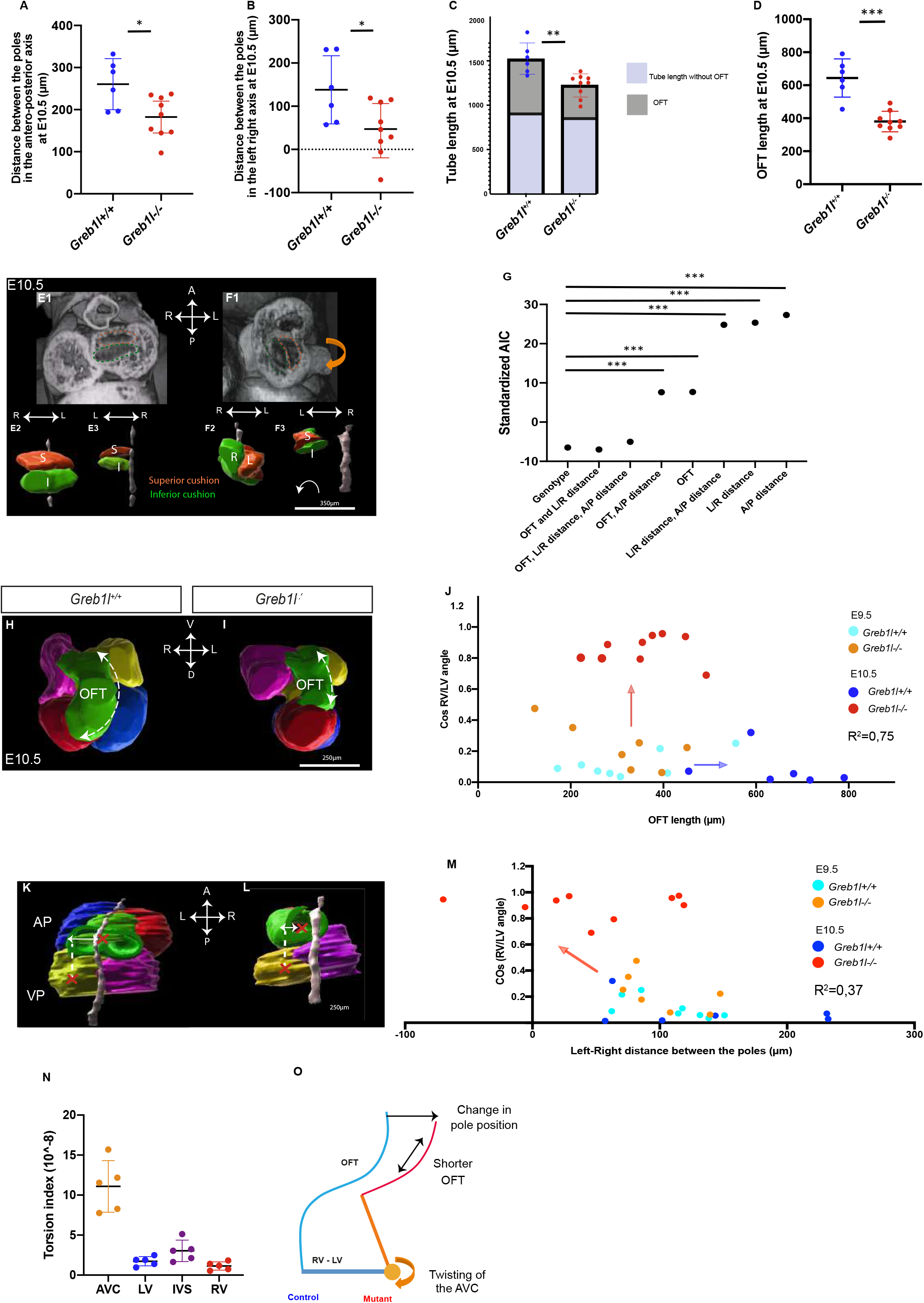
**Outflow tract and atrioventricular canal defects in *Greb1l* mutants at E10.5.** (A-B) Quantification of the antero-posterior (A) and left-right (B) distance between the heart tube poles in controls (n=8) and mutants (n=9) at E10.5. *p-value<0.05 (t-test). (C-D) Quantification of the heart tube length at E10.5 in the same samples, showing the contribution of the outflow tract (OFT, grey in C, and D) relative to the rest of the tube (pale blue in C). **p<0.01 (t-test). (E-F) 3D images by HREM showing coronal sections of atrioventricular cushions in controls (E) and *Greb1l* mutants (F) and associated 3D segmentations in ventral (E2-F2) and dorsal (E3-F3) views. Superior (S) and inferior (I) cushions (proximally) are in orange and green respectively, the notochord in grey. The orientation of cushion spiralling is indicated by an arrow. (G) Values of the standardised Akaike Information Criterion (AIC) for the indicated combination of parameters in a Generalized Linear Model. ***relative likelihood<0.001 (n=15). (H-I) 3D segmented heart at E10.5 in a cranial view. Arrows outline differential lengths of the outflow tract in controls and mutants. (J) Correlation between the outflow tract length (x) and the RV/LV axis (y), shown at E9.5 (paler colours) and E10.5 (darker colours) in controls (blue) and mutants (orange and red). Significant changes between stages are indicated by arrows. R^2^, Pearson determination coefficient at E10.5 (n= 7 and 6 controls at E9.5 and E10.5, 7 and 9 mutants at E9.5 and E10.5, respectively) (K-L) 3D segmented heart at E10.5 in a dorsal view. Arrows indicate differential left-right distances between the poles (red crosses) in controls and mutants. (M) Correlation between the left-right pole distance (x) and the RV/LV axis (y) shown as in J. (N) Torsion index calculated for heart tube regions. Mean values and standard deviations are shown for 5 control hearts at E10.5. (O) Schematic representation of the mechanical model, whereby twisting (arrow) of the AVC results from growth arrest in the OFT and reduced pole distance. The control and mutant situations are in blue and orange respectively. Means and standard deviations are shown. A, anterior; AP, arterial pole; AVC, atrioventricular canal; D, dorsal; IVS, interventricular septum; L, left; LV, left ventricle; R, right; RV, right ventricle; OFT, outflow tract; P, posterior; V, ventral; VP, venous pole.

To understand the mechanism of abnormal ventricle position, we investigated which geometrical parameters were more closely associated with ventricle position, using a Generalized Linear Model. The standardised Akaike Information Criterion (AIC) of the model was best when we considered both the outflow tract length and the left-right distance between the heart tube poles (Fig. 4G), and indeed was very close to the AIC of a model using the genotype as variable. We then analysed the kinetics of these parameters in relation to ventricle position. Between E9.5 and E10.5, whereas the outflow tract grows 2 fold in control hearts (325±1123µm in n=8 E9.5 compared to 644±116µm in n= 6 E10.5, p<0.0001, t-test), growth was completely arrested in *Greb1l* mutants (309±112µm in 7 E9.5 compared to 380±62µm in 9 E10.5, p=0.13, t-test), the length of the OFT at E10.5 being a strong linear predictor (R^2^=0.75) of the ventricle position (Fig. 4H-J). During the same time window, the left-right distance between the heart tube poles, which tends to be stable in control hearts, was significantly reduced in *Greb1l* mutants, this distance being also a linear predictor of the ventricle position, although to a lesser extent (R^2^=0.37) (Fig. 4K-M).

Our quantitative anatomical analyses, combined with the expression of *Greb1l* mainly outside the heart tube, and with normal heart tube patterning in the mutant, suggest a mechanical process of ventricle malpositioning. We propose a model in which differential growth in the outflow tract relative to the rest of the heart tube, as well as reduction of pole distance, exert a constraint which pulls the tube towards the arterial pole. Since the atrioventricular canal is the section of the heart tube with the least resistance to torsion (Fig. 4N), pulling would result in twisting at the atrioventricular canal (Fig. 4O).

### GREB1L is a novel player in retinoic acid signalling

*Greb1l* has been shown previously to be a direct target of RA receptors (Simandi et al., 2016) and RA signalling is required for outflow tract growth (Duong et al., 2021; El Robrini et al., 2016; Li et al., 2010). These led us to hypothesise that *Greb1l* could interfere with RA signalling. We first analysed the expression pattern of *Greb1l* relative to that of *Aldh1a2*, encoding the main RA synthesising enzyme, and of the transgenic reporter *RARE-lacZ*. In the heart field, *Aldh1a2* was expressed in the most posterior third, overlapping with *Greb1l*, which covers the whole heart field (Fig. 5B). *RARE-lacZ* extended more anteriorly over two thirds, also overlapping with *Greb1l* (Fig. 5A). This indicates that *Greb1l* is expressed within the RA responsive domain, but also extends more broadly. Genetic tracing with either *RARE-lacZ* or *RARE-CreERT2;R26^mTmG/+^* shows that RA responsive cells are fated to the outflow tract myocardium at E8.5j-E9.5, in addition to that of the atria (Fig. S2A-D).

**Figure 5.**
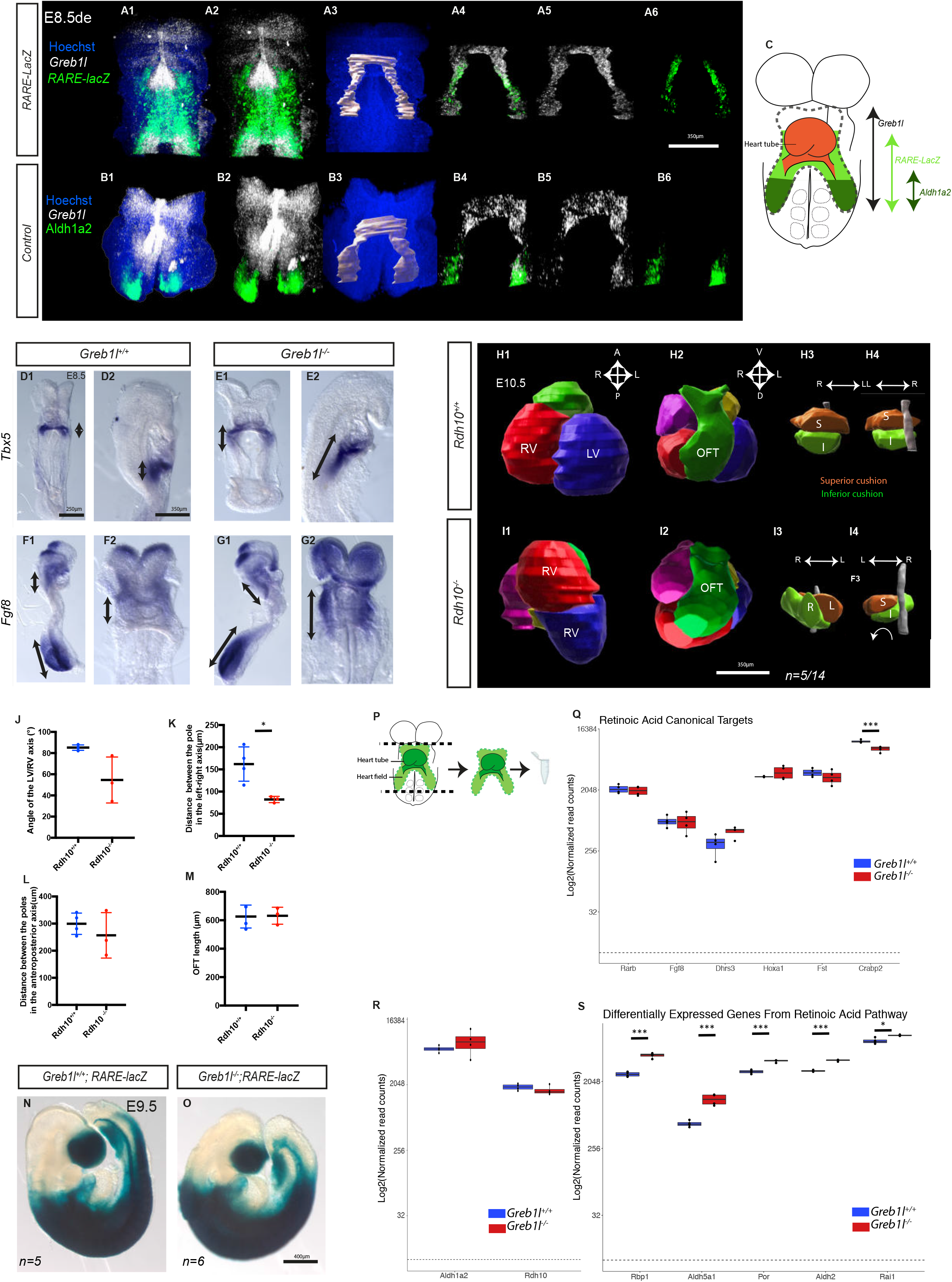
***Greb1l* mutants phenocopy partial deficiency in retinoic acid.** (A-B) *Greb1l* (white), *lacZ* (green, A) and *Aldh1a2* (green, B) expression at E8.5de detected by double whole mount RNA scope ISH in transgenic *RARE-lacZ* (A) and control (B) embryos, shown in frontal views. After segmentation (A3-B3), staining in the heart field was extracted (A4-B6). (C) Schema recapitulating the overlap of gene expression in the heart field. (D-G) *Tbx5* (D-E) and *Fgf8* (F-G) expression in control (n=5) and *Greb1l^ex17/ex17^* mutants (n=4) at E.8.5, shown in ventral (D1, E1, F2, G2) and right-sided (D2, E2, F1, G1) views. Black double arrows indicate differential expansion of the staining along the antero-posterior axis. (H-I) 3D segmentation of control and *Rdh10* mutant hearts at E10.5, in frontal (H1-J1) and cranial (H2-J2) views. In H3-I3, the superior (S) and inferior (I) cushions (proximally) are segmented in orange and green respectively, the notochord in grey. The orientation of cushion spiralling is indicated by an arrow. (J-M) Corresponding quantifications of the RV/LV axis (J), left-right (K) and antero-posterior (L) distance between the heart tube poles, and outflow tract length (M) in controls (n=3) and *Rdh10* mutants (n=3) at E10.5. * p-value<0.05 (Mann Whitney test in J, t-test in K-M). (N-O) Brightfield images of *Greb1l^+/+^;RARE-lacZ* control and *Greb1l^-/-^;RARE-lacZ* mutants at E9.5. β-galactosidase staining is in blue. The dotted yellow line represents the anterior limit of the staining. (P) Outline (green) of the region isolated for RNA-sequencing at E8.5e. (Q) Normalized read counts of canonical target genes of RA signalling, with functional RARE sequences. ***FDR<0.001 (n=4 controls, 4 *Greb1l^-/-^* mutants, DESeq2). (R-S) Normalized read counts of genes involved in the RA pathway, not (R) or differentially expressed (S) between controls (blue) and *Greb1l^-/-^* mutants (red). The dotted line indicates the threshold of background expression. *FDR<0.05, ***FDR<0.001 (n=4 controls, 4 *Greb1l^-/-^* mutants, DESeq2). Means and standard deviations are shown. A, anterior; D, dorsal; FDR, false discovery rate; IVS, interventricular septum; L, left; LV, left ventricle; R, right; RV, right ventricle; OFT, outflow tract; P, posterior; V, ventral. See also Figure S2.

We then investigated whether RA signalling is affected in *Greb1l* mutants, using canonical transcriptional readouts. Abnormal patterning of rhombomeres in *Greb1l* mutants (Fig. S2F-G), in which rhombomere 4 is present but posteriorized, is a characteristic signature of partial RA deficiency seen in *Hnf1b* and *Rdh10* mutants (Rhinn et al., 2011; Sirbu et al., 2005), in contrast to the loss of r4 upon complete RA deficiency in *Aldh1a2* mutants (Niederreither et al., 2000). Within the cardiac region, the expression profile of two RA responsive genes, *Tbx5* and *Fgf8* (Cunningham et al., 2013; Niederreither et al., 2001; Ryckebusch et al., 2008; Sirbu et al., 2008), were expanded posteriorly in *Greb1l* mutants compared to control embryos (Fig. 5D-G). This abnormal molecular patterning phenocopies RA deficiency in *Aldh1a2* and *Rdh10* mutant embryos (Cunningham et al., 2013; Niederreither et al., 2001; Sirbu et al., 2008). At a morphological level, complete RA deficiency in *Aldh1a2* mutants is more severe than *Greb1l* mutants : earlier lethality (E10.5), early defects in heart tube formation (Niederreither et al., 2001; Ryckebusch et al., 2008). Thus, we focussed on a model of partial RA deficiency, in which another RA synthesising enzyme, Rdh10, is targeted. *Rdh10* mutants were reported to survive until E12.5 with a milder phenotype of “misaligned putative ventricles” (Rhinn et al., 2011). We analysed this into more detail. At E10.5, 5/14 (35%) *Rdh10*^-/-^ embryos had abnormal supero-inferior ventricles and spiral atrioventricular cushions indicative of criss-cross heart, with variable severity (Fig. 5H-J). This was associated with a reduction in the left-right pole distance, but not with a shorter outflow tract (Fig. 5K-M). Thus, our analyses show that partial RA deficiency in *Rdh10* mutants leads to criss-cross heart with supero-inferior ventricles and reduction in the left-right pole distance. Molecular patterning and morphological observations both support striking similarities between *Greb1l* inactivation and partial RA deficiency.

To further monitor RA response, we analysed the expression of the transgenic reporter *RARE-lacZ* in *Greb1l* mutants. This was largely unaffected in *Greb1l* mutants, with a similar anterior boundary of expression (Fig. 5N-O). The less posterior boundary is in keeping with the phenotypic posterior truncation. The maintenance of *RARE-lacZ* in *Greb1l* mutants thus differs from its absence in *Aldh1a2* and *Rdh10* mutants (Cunningham et al., 2011; Niederreither et al., 1999). We next performed a transcriptomic profiling of the cardiac region in *Greb1l* mutant compared to control embryos at E8.5e (Fig. 5P), at the end of the time-window of *Greb1l* expression in controls. After quality check (Fig. S3A-C), we identified significantly differentially expressed genes in both *Greb1l* mutant lines. In agreement with the *RARE-lacZ* profile, canonical RA targets with functional retinoic acid responsive elements (RARE) (Berenguer et al., 2020), including *Rarb*, *Fgf8*, *Dhrs3*, *Hoxa1* and *Fst*, had normal expression levels in *Greb1l* mutants, except *Crabp2*, which was significantly downregulated (Fig. 5Q). In addition, expression of *Aldh1a2* and *Rdh10*, which encode the main RA synthesising enzymes in cardiac cells, were unchanged (Fig. 5R). However, other components of the RA pathway were significantly upregulated, including *Aldh2, Aldh5a1*, *Por*, *Rai1 and Rbp1* (Fig. 5S), which is compatible with compensatory mechanisms, as shown previously (D’Aniello et al., 2013), of a phenotype equivalent to partial RA deficiency. This suggests that GREB1L does not fundamentally affect canonical RA signalling via RARE sequences.

Taken together, our results show that Greb1l inactivation phenocopies the specific features of partial RA deficiency, but without affecting canonical RA signalling via RARE sequences. This suggests that GREB1L is a novel modulator of RA signalling.

### *Greb1l* regulates cardiomyocyte differentiation and ribosome biogenesis to maintain the pool of precursor cells

Given that *Greb1l* is expressed more broadly than the RA responsive domain, we analysed its role independently of RA signalling, using transcriptomic approaches. We identified 293 genes consistently deregulated in both *Greb1l* mutant lines (Fig. 6A-B) and selected four genes, *Cacna1h*, *Fabp3*, *Col5a1* and *Hapln1* to validate differential expression (Fig. 6G-J, Fig. S3D-H). Then, we intersected differentially expressed genes with the list of genes significantly correlated with variations in *Greb1l* expression levels in single wild-type embryonic cardiac cells (Fig. 6B-C). Satisfactorily, 88 genes negatively correlated with *Greb1l* in wild-type cells, were significantly upregulated in *Greb1l* mutants, whereas 161 genes positively correlated with *Greb1l* in wild-type cells were significantly downregulated in *Greb1l* mutants (Fig. 6C). We have thus identified a set of genes specifically associated with *Greb1l* in embryonic cardiac cells (Table 1). Enrichment analysis of GO pathways singles out two main categories of genes : genes involved in cardiomyocyte differentiation are collectively enriched in *Greb1l* mutants, whereas genes associated with ribosomes are collectively depleted in *Greb1l* mutants (Fig. 6D). We further analysed targeted gene lists of cardiomyocyte differentiation and ribosomes. This shows that genes specifically involved in terminal cardiomyocyte differentiation (Me3) rather than intermediate cardiomyocyte differentiation (Me4/6) are upregulated in *Greb1l* mutants (Fig. 6E, S3I). In addition, genes specifically involved in ribosome biogenesis, rather than ribosome components (Fig. 6F, S3J) are downregulated in *Greb1l* mutants. *Ncl*, which encodes a major rRNA processing protein (Ginisty et al., 1998), is the most significant gene of the list (correlation FDR=6.7x10^-15^, differential expression FDR=8.4x10^-22^, Table1).

**Figure 6.**
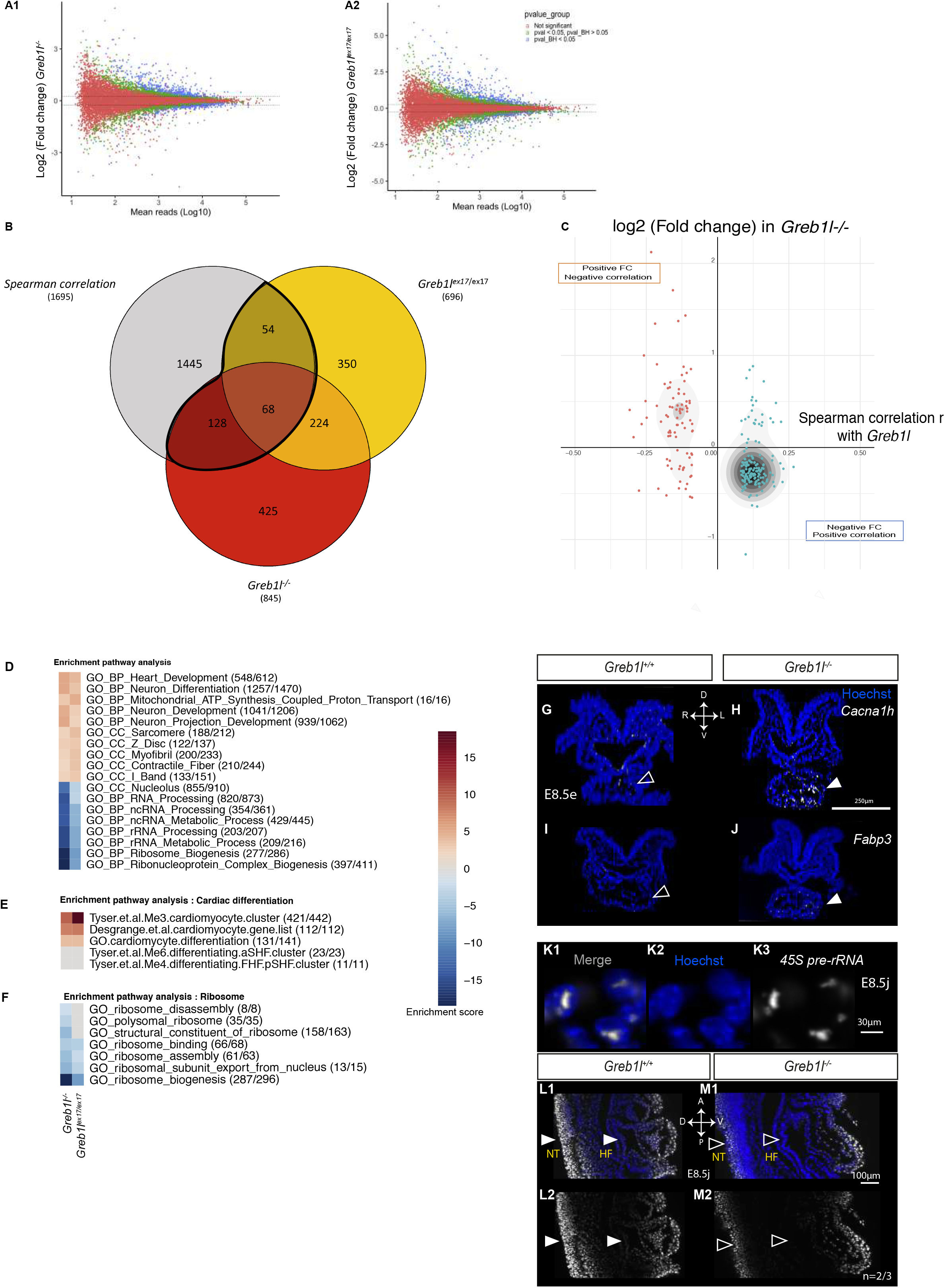
**Transcriptomic changes in cardiomyocyte differentiation and ribosome biogenesis downstream of *Greb1l*.** (A) MA plots representing relative gene expression between littermate controls and *Greb1l^-/-^* (A1) or *Greb1l^ex17/ex17^* (A2) mutants at E8.5e. Non-significant differential expression is represented in red, differential expression in green (p-value<0.05), and blue (Benjamini-Hochberg (BH) corrected p-value<0.05, DEseq2, n=4 controls, 4 *Greb1l^-/-^* mutants, 4 *Greb1l^ex17/ex17^* mutants). (B) Venn diagram comparing differentially expressed genes in *Greb1l^ex17/ex17^* (yellow) and *Greb1l^-/-^* (red) mutants with genes correlated with *Greb1l* levels in single wild-type embryonic cardiac cells (gray, see also Fig. 2G). The 249 genes (excluding *Greb1l*) used for further analysis are outlined in bold. (C) Corresponding representation of the 249 genes, according to their fold change (FC) in *Greb1l^-/-^* mutants and their Spearman correlation coefficient (red, r<0; blue, r>0, see Fig. 2G). Point density is represented by a gray scale. Genes positively correlated with *Greb1l* in wild-type cells are significantly downregulated in mutants, and vice-versa. (n=249, p<0.0001, chi2 test). (D) Heatmap representing the most enriched GO pathways, ordered according to their enrichment score (blue, negative; gray, null; red, positive) in *Greb1l^-/-^* (left column) and *Greb1l^ex17/ex17^* (right column) mutants. Numbers in parentheses indicate the fraction of genes considered in specific GO terms. (E-F) Heatmaps of gene set enrichment associated with cardiac differentiation (E) and ribosome (F), represented as in D. See gene lists in (Desgrange et al., 2020; Tyser et al., 2021). (G-J) Expression of cardiomyocyte genes *Cacna1h* (G-H) and *Fabp3* (I-J) in controls (G, I, n=3 each) and *Greb1l^-/-^* mutants (H, J, n=3 each) detected by whole mount RNAscope ISH at E8.5e, and shown in transverse sections. Filled and empty arrowheads point to high and low expression in the heart, respectively. (K-M) Expression of the 45S pre-ribosomal rRNA detected by whole mount RNAscope ISH in controls (K-L, n=3 from the *Greb1l^ex17^* line) and *Greb1l^-/-^* mutants (M) at E8.5j. A, anterior; D, dorsal; FDR, false discovery rate; HF, heart field; L, left; LV, left ventricle; NT, neural tube; R, right; RV, right ventricle; OFT, outflow tract; P, posterior; V, ventral. See also Table 1 and Figure S3.

**Table 1:** List of 249 *Greb1l* dependent genes. Genes shown have been filtered for an FDR<0.05 in the Spearman correlation with *Greb1l* levels in single cell transcriptomes of wild-type embryonic cardiac cells (columns D-E). Genes with r>0 and r<0 are shown in separate sheets. Additionally, genes have been filtered for an FDR<0.05 in the differential expression between *Greb1l* mutants and controls using the DEseq2 package (columns F-I).

To validate pathway analysis, we performed in vivo labelling experiments. Ribosome biogenesis (Hadjiolov and Nikolaev, 1976) was evaluated by in situ hybridisation of the 45S pre-ribosomal rRNA at E8.5j, a few hours after we have seen ribosome gene downregulation. Nucleolar staining of 45S pre-rRNA was decreased in the heart field and neural tube, but not in the heart tube, reflecting *Greb1l* expression domains (Fig. 6K-M). Cardiomyocyte differentiation genes such as *Cacna1h* and *Fabp3* were more highly expressed in *Greb1l* mutant hearts at E8.5e, indicating premature differentiation in accordance with transcriptomic analysis at the same stage (Fig. 6G-J). At a later stage, E9.5, a few cardiomyocytes were abnormally detected in the dorsal pericardial wall of *Greb1l* mutants (Fig. 7A1-B1). More strikingly, the heart field, labelled by Isl1, was found significantly reduced in size and disorganised in *Greb1l* mutants (Fig. 7A2-B4, C), indicative of an exhaustion of the precursor cell reservoir. In addition, the cardiac precursor marker Isl1 was found more inside the heart tube, in the right atrium (Fig. 7A3-B3). Conversely, a more mature cardiomyocyte gene, *Fabp3*, had a reduced expression at the heart poles (Fig. 7D-E, Fig. S3K), specifically in the inferior outflow tract and dorsal right atrium that are labelled by *RARE-lacZ* (Fig. 7F-G). The defects in cardiomyocyte differentiation that we detect at the poles of the heart tube, where cells ingress to elongate the tube, prefigure growth arrest in the outflow tract at E10.5.

**Figure 7.**
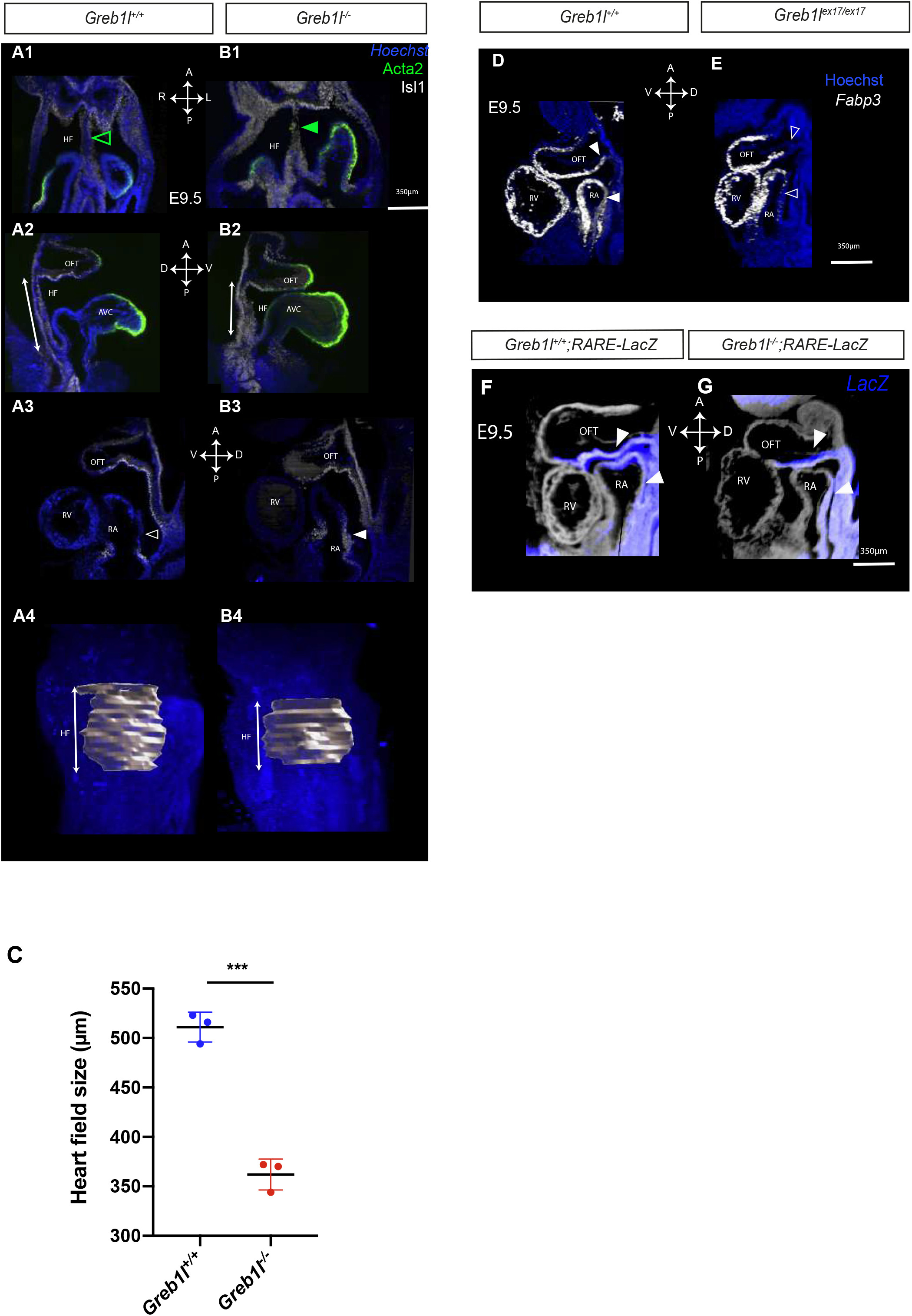
**Defective heart field size and cell differentiation at both poles of *Greb1l* mutant hearts at E9.5.** (A-B) Expression of cardiomyocyte (Acta2) and heart field (Isl1) markers in controls (A, n=3) and *Greb1l^-/-^* mutants (B, n=3) detected by whole mount immunofluorescence at E9.5, and shown in frontal (A1-B1) and sagittal (A2-B3) sections. The double arrow points to the size of the heart field in a midline sagittal section. 3D segmentation of the heart field is shown in A4-B4. (C) Corresponding quantification of heart field size. ***p<0.001 (t-test). (D-E) Expression of the cardiomyocyte gene *Fabp3* in controls (D, n=5) and *Greb1l^ex17/ex17^* mutants (E, n=5) detected by whole mount RNAscope ISH at E9.5, and shown in sagittal sections. Filled and empty arrowheads point to high and low expression in cardiac regions. (F-G) HREM sagittal sections of *Greb1l^+/+^;RARE-lacZ* controls and *Greb1l^-/-^;RARE-lacZ* mutants at E9.5 stained for β-galactosidase activity (blue). Means and standard deviations are shown. A, anterior; AVC, atrioventricular canal; D, dorsal; HF, heart field; L, left; LV, left ventricle; NT, neural tube; R, right; RA, right atrium; RV, right ventricle; OFT, outflow tract; P, posterior; V, ventral.

Taken together, our transcriptomic analyses and validations provide the first demonstration of a function of *Greb1l* in promoting ribosome biogenesis and controlling cell differentiation, to preserve the pool of precursor cells required for the elongation of the heart tube.

## Discussion

We have uncovered the role of *Greb1l* in maintaining a reservoir of precursor cells, which supplies the heart tube and shapes it. With quantitative spatio-temporal gene expression analyses, we show that *Greb1l* is expressed transiently in early cardiac precursors at E8.5d-f. It promotes ribosome biogenesis and controls cell differentiation, intervene in RA signalling. The absence of *Greb1l* prevents the elongation of the heart tube after E9.5 and reduces the distance between the tube poles. This is predictive of heart tube remodelling after heart looping, including torsion of the atrioventricular canal and supero-inferior ventricles. By characterizing the first mouse model of criss-cross heart, our work thus identifies the embryological mechanisms for this congenital heart defect.

Heart morphogenesis relies on the formation and shaping of an embryonic tube. The rightward looping of the tube is important for positioning the right ventricle and thus the overall alignment of cardiac chambers (Desgrange et al., 2020; Le Garrec et al., 2017; Patten, 1922). Our analysis of *Greb1l* mutants now indicates that the position of ventricles is not fixed after heart looping, and that later growth of the heart tube can further modulate ventricle position. We thus uncover a novel aspect of heart morphogenesis, corresponding to the maintenance of chamber position subsequent to heart looping. Heart looping is determined by mechanical constraints during tube growth, such that tube deformation is primarily driven by a buckling mechanism (Le Garrec et al., 2017; Patten, 1922). Similarly, for the maintenance of chamber position, we find that growth of the heart tube, by ingression of precursor cells from the heart field, is essential. Impairment of growth in the absence of *Greb1l* generates abnormal mechanical constraints, which remodel the heart tube after heart looping. Another parameter of tube remodelling both during looping (Le Garrec et al., 2017) and thereafter as shown here, is the distance between the poles. This is the main parameter associated with supero-inferior ventricles in *Rdh10* mutants. However, supero-inferior ventricles are fully penetrant, when both pole distance and outflow tract growth are affected in *Greb1l* mutants. The stages after heart looping have been previously described as a convergence step, in keeping with the displacement of the venous pole, from a caudal location at E8.5, to dorsal to the outflow tract at E10.5 (Patten, 1922). However, our quantifications of pole distance along the antero-posterior axis rather indicate a significant increase between E9.5 and E10.5, from 112μm±40 to 260μm±57 (p<0.001, t-test, Fig. 3H, 4A). This indicates that there is no measurable convergence, which is thus an illusion: from the growth of the atria and outflow tract, which appear closer, and from the growth of the ventricles which protrude ventrally and are displaced caudal to the atria.

Abnormal ventricle position is classically associated with the heterotaxy syndrome (Jacobs et al., 2007; Van Praagh, 1972), which results from impaired left-right signalling (Hummel and Chapman, 1959; Meno et al., 1998; Supp et al., 1997). However, criss-cross heart is anatomically different from heterotaxy, as primarily defined by crossing of atrioventricular connections (Anderson, 1982; Manuel et al., 2018). Our results highlight distinct morphogenetic and molecular mechanisms for ventricle malposition in criss-cross heart, compared to heterotaxy. Whereas heterotaxy is associated with defective heart looping (Campione et al., 1999; De La Cruz et al., 1959; Desgrange et al., 2020; Van Praagh et al., 1964; Yan et al., 1999), we show that criss-cross heart is not. In addition, heterotaxy results from impaired left-right signalling upon mutations in ciliary genes or in Nodal pathway components (Hummel and Chapman, 1959; Meno et al., 1998; Supp et al., 1997) whereas we identify a novel gene network associated with criss-cross heart, involving *Greb1l*.

Our quantification of shape changes in a unique mouse model uncovers the pathological mechanisms for criss-cross heart. This congenital heart defect is thus primarily a shortage of precursor cells, which will most strikingly arrest the growth of the outflow tract. This explains why criss-cross heart is frequently associated with malposition of the great arteries, pulmonary stenosis and right ventricle hypoplasia (Freedom et al., 1978; Manuel et al., 2018). Hypoplasia is thus not an indirect effect of heart rotation as hypothesised previously (Seo et al., 1992). As a consequence of growth anomalies and reduction in pole distance, the embryonic heart tube is remodelled. This includes a torsion of the tube at the level of the atrioventricular canal, which accounts for twisted atrioventricular connections in criss-cross heart (Seo et al., 1992). Our observations at sequential stages of development indicate that the timing of twisting is before heart septation, between E9.5 and E10.5, which differs from predictions by anatomists (Anderson, 1982; Anderson et al., 1974). Thus, our analyses of *Greb1l* mutants provide a comprehensive mechanistic insight into criss-cross heart and associated cardiac defects. However, criss-cross heart is not always associated with supero-inferior ventricles (Manuel et al., 2018). This may reflect different degree of tube torsion (Freedom et al., 1978), or raise the possibility that ventricle position may be corrected by further growth of the heart after E10.5.

Elongation of the heart tube depends on the progressive incorporation of precursor cells from the dorsal pericardial wall, pharyngeal mesoderm and juxta-cardiac field abutting the septum transversum (Domínguez et al., 2012; Kelly et al., 2001; Tyser et al., 2021). Impairment of cardiomyocyte differentiation in *Nkx2-5*, *Tbx20, Mef2c* or *Asb2* mutants, arrests heart development at E8.5-E9.5 and is embryonic lethal (Lin et al., 1997; Lyons et al., 1995; Métais et al., 2018; Stennard et al., 2005). Decreased proliferation of heart precursors in *Fgf8* or *Hes1* mutants (Park et al., 2006; Rochais et al., 2009), or disruption of the second heart field epithelial architecture in *Vangl2* and *Tbx1* mutants, which induces cardiomyocyte differentiation (Francou et al., 2014; Ramsbottom et al., 2014) shorten the outflow tract. These mutants, which survive to birth, manifest conotruncal malformations and ventricular septal defects. Anomalies in ventricle position seem to be visible at E9.5-E10.5 but have not been reported at birth, and neither criss-cross heart. This indicates that impaired outflow tract growth not always results in abnormal remodelling of the cardiac tube. It remains to be investigated whether this is due to a threshold in the intensity of growth defects or in the specific timing of growth defects, since *Greb1l* is transiently expressed in heart precursors at E8.5c-f.

RA, which is known to regulate cardiomyocyte differentiation, heart field size (Duong et al., 2021; Keegan et al., 2005; Ryckebusch et al., 2008) and outflow tract growth (Duong et al., 2021; El Robrini et al., 2016; Li et al., 2010), had been previously identified in ES cells as an inducer of *Greb1l* expression, via a RARE sequence 13kb upstream of *Greb1l* promoter (Simandi et al., 2016). We now show in vivo that partial RA deficiency phenocopies *Greb1l* inactivation : *Rdh10* mutants have criss-cross heart with supero-inferior ventricles and other models of transient RA deficiency, *CAGG-Cre-Esr1;RARa1^-/-^, RXRa^flox/flox^* conditional mutants or RA-rescued *Aldh1a2* mutants, display outflow tract shortening (El Robrini et al., 2016; Li et al., 2010). This functional evidence would position GREB1L as a modulator of RA signaling, although its mechanism of action remains unknown. The observation that *Greb1l* expression extends more anteriorly than the RA-responsive domain suggests that *Greb1l* expression integrates other upstream regulators. Whereas RA signalling has been shown to regulate gene expression by the binding of RA receptors to RARE sequences in target genes, we found in *Greb1l* mutants a normal pattern of the *RARE-lacZ* reporter transgene. In keeping with this, ChIP-seq and transcriptomic analyses in the paraxial mesoderm have shown that 50% of retinoic acid targets are devoid of RARE sequences (Berenguer et al., 2020). Thus, GREB1L is a good candidate for taking part in alternative transcriptional complexes interacting with RA signalling to regulate gene expression. Proteomics and ChIP-seq analyses will be important to further test this hypothesis. By cross-correlated transcriptomic analyses, we have identified a gene set specifically associated with *Greb1l*.

Our work on a novel marker of heart precursors, *Greb1l*, reveals a novel dimension in the regulation of the reservoir of precursor cells. In addition to precocious cardiomyocyte differentiation, transcriptomics of *Greb1l* mutants reveals reduced ribosome biogenesis. Although ribosomes are essential in every cell for protein synthesis, congenital ribosomopathies have tissue-specific phenotypes (Luzzatto and Karadimitris, 1998; Orgebin et al., 2020), including conotruncal defects in Diamond Blackfan Anemia (Vlachos et al., 2018). Ribosome levels have been shown to vary along cell trajectories, being lower in quiescent and differentiated cells but higher in intermediate cycling precursors (Chau et al.; van Velthoven and Rando, 2019), and for example higher in human ES cells compared to derived cardiomyocytes (Pereira et al., 2019). In agreement with these data, we show that impaired ribosome biogenesis in *Greb1l* mutants correlates with impaired cardiomyocyte differentiation and an overall growth arrest of the outflow tract. The reduced size and disorganisation of the heart field further show the exhaustion of the reservoir of cardiac precursor cells in *Greb1l* mutants. This may occur not only by cell differentiation but potentially also by mechanisms common to ribosomopathies, such as cellular hypo-proliferation, proteolytic and oxidative stress (Kampen et al., 2020).

*GREB1L* has been previously associated with a broad spectrum of congenital defects in humans, such as urogenital defects, hearing impairment and skeletal anomalies (Schrauwen et al., 2020). *Greb1l* mouse mutants are more severe and display general growth retardation, neural tube defects and craniofacial anomalies (De Tomasi et al., 2017 and Fig. S1). All these phenotypes are also observed in ribosomopathies (Orgebin et al., 2020), consistently with the role of *Greb1l* in regulating ribosome biogenesis. Despite the number of genetic variants in patient cohorts, *GREB1L* has remained a poorly characterized gene. Bioinformatics predictions (Uniprot, MobiDB, Necci et al., 2017) indicate the presence of intrinsically disordered regions in GREB1L, which is compatible with a nucleolar localization (Guillen-Chable et al., 2021), where ribosomes are produced. Alternatively, by analogy with its paralogue GREB1, GREB1L could be involved in protein O-GlcNAcylation (Shin et al., 2021), which is a modification required for the stability of ribonucleoproteins (Qin et al., 2017). The processes regulated by *Greb1l* that we have discovered during cardiac development thus provide novel insight into a broad spectrum of congenital defects.

## Acknowledgments

We thank V. Benhamo, L. Guillemot, M. Cavaignac, M. Franco and C. Cimper for generous technical assistance, T. Holm Bønnelykke for expert advice, A. Schedl, V. Ribes, S. Zaffran, E. Calo, M. Cohen-Tannoudji for insightful discussions, A. Schedl, G. Comai, V. Fraulob for the production of *Tg(RARE/Hspa1b-cre/ERT2)#Asc;R26^mTmG/mTmG^* and *Rdh10* mutant embryos, C. Jeanpierre and L. de Tomasi for *Greb1l* mutant mice, D. Conrozet and the Histology platform of the SFR Necker, S. Dupichaud and the Imaging platform, N. Goudin and the Image Analysis plateform, C. Bole-Feysot and M. Zarhrate of the Genomics platform, N. Cagnard and the Bioinformatics platform, Y. Zimmermann and the LEAT animal facility. This work was supported by core funding from the Institut Pasteur and INSERM, state fundings from the Agence Nationale de la Recherche under ‘‘Investissements d’avenir’’ program (ANR-10-IAHU-01, ANR-10-LABX-73-01 REVIVE), grants from the ANR (ANR-21-CE14-0062-01) and the Fondation Française de Cardiologie to S.M.M., the MSD-Avenir fund (Devo-Decode project) and the AXA Research Fund. S.B. has benefited from an MD-PhD fellowship of the Institut *Imagine*, supported by State funding from the ANR under “Investissements d’avenir” program (ANR-10-IAHU-01) and the Fondation Bettencourt-Schueller, and training from the Ecole Doctorale FIRE - Programme Bettencourt-Schueller. A.B., F.R., D.B., L.H. are supported by the APHP. S.M.M. is an INSERM research director.

## Author contributions

Conceptualization, S.B., S.M.M.; Methodology, S.B., J.-F.LG., E.P. and S.M.M.; Software, J.-F.LG. and E.P.; Formal Analysis, S.B., A.B., E.P. and L.H.; Investigation, S.B., A.B. and A.D.; Resources : W.K., L.H., F.R. and D.B.; Writing – Original Draft, S.M.M.; Writing – Review & Editing, all authors; Visualization, S.B., A.B., and E.P.; Supervision, S.M.M. and S.B.; Project Administration, S.M.M.; Funding Acquisition, S.M.M., S.B., A.B.

## STAR Methods

### EXPERIMENTAL MODEL AND SUBJECT DETAILS

#### Animals

Wild-type mouse embryos were from a C57Bl6J genetic background. *Tg(RARE-Hspa1b/lacZ)12Jrt/J* (Rossant et al., 1991), *Tg(RARE/Hspa1b-cre/ERT2)#Asc* (Da Silva et al., 2021), *Rdh10^tm1.1Ics^* (Rhinn et al., 2011), Gt(ROSA)26Sortm4(ACTB-tdTomato,-EGFP)Luo (referred to as *R26^mTmG^*, Muzumdar et al., 2007) mice were maintained in a mixed genetic background. The *Greb1l^em1Cjea^* (referred to as *Grebl1^-/-^* throughout the paper) and *Greb1l^ex17^* mutant lines were generated in a C57BL6J genetic background (De Tomasi et al., 2017). By Sanger sequencing, we characterized a 5bp deletion, c.2403_2407delCGGAT, in the *Greb1l^ex17^* line used here. Embryonic day (E) 0.5 was defined as noon on the day of vaginal plug detection. Heart looping stages from E8.5c to E8.5j were defined according to the previously published nomenclature (Le Garrec et al., 2017). Both male and female embryos were collected and used randomly for experiments. Control embryos are wild-type littermates, unless specified. All embryos were genotyped by PCR (378bp and 757pb amplicon for the *Greb1l^em1Cjea^* and *Greb1l^ex17^* alleles respectively), using distinct primer pairs for the wild-type and deleted alleles to detect the small deletion. *Greb1l^-/-^* mutants die between E14.5 and P0. *Greb1l^ex17/ex17^* die at E11.5 and present a growth delay from E9.5. Animals were housed in the Laboratory of Animal Experimentation and Transgenesis of the SFR Necker, Imagine Campus, Paris, and in the animal facility of the Institut Pasteur. Animal procedures were approved by the ethical committees of the Institut Pasteur and Université de Paris and by the French Ministry of Research.

#### Patient recruitment and cardiac imaging

3D images of human hearts were selected from available cardiac CT-scans of patients with criss-cross hearts and control patients in the department of Pediatric Radiology in Necker Hospital. The CT scan images were acquired and completely anonymized, in accordance with the procedures of the APHP data protection office (N° 20201112163629). Patient CT examinations were done using a Lightspeed VCT 64-slices, GE Healthcare (Coverage 4 cm of patient anatomy per rotation, gathering 64 slices at 0.625 mm)(Habib Geryes et al., 2016). During CT acquisitions, contrast (Iomeron® 300 mg/ml, 2mL/kg) was injected at a flow rate determined by body size and intravenous access size (1.5 ml/s up to 2 ml/s) followed by a saline flush using a power injector. Image reconstruction was performed with a slice thickness of 0.625 mm, an increment of 0.625 mm, and the STANDARD reconstruction kernel. Iterative reconstruction was used with 60 % ASIR.

ECG-gated scans were performed in SAS (Step And Shoot) mode at a single phase (75 %) in diastole for a heart rate < 65 bpm and a single phase (40 %) in systole for heart rate ≥ 65 bpm. In patients with heart rate < 65 bpm, reconstructions were performed at mid-diastole and in patients with heart rate ≥ 65 bpm, reconstructions were performed at end-systole. SAS scans were done with detector collimation of 64 × 0.625 mm, a gantry rotation time of 350 ms. Non-gated scans for the examination of the whole thorax were performed with detector collimation of 64 × 0.625 mm, a gantry rotation time of 400 ms. A 0.984 pitch factor, 80 kV tube voltage and modulated mA tube current between 100 and 230 mA with a noise index of 25 Hounsfield Unit were used. Automatic segmentation and 3D measures were made using the Imaris Software.

### METHOD DETAILS

#### β-galactosidase staining

Embryos were collected at E9.5. The heart was arrested in diastole with 250mM cold KCl (E9.5). *Tg(RARE-Hspa1b/lacZ)12Jrt/J* embryos were fixed in 4% PFA – 5mM EGTA – 2mM MgCl2 for 10min. Embryos where then permeabilized in 0.2% NP40 – 2mM MgCl2 – 0.1% sodium deoxycholate 30min and stained wholemount overnight in Xgal solution. Brightfield images were acquired with a Zeiss AxioCamICc5 Camera and a Zeiss StereoDiscovery V20 stereomicroscope with a Plan Apo 1.0X objective.

#### **RNA *in situ* hybridisation**

ISH was performed on wholemount embryos at E8.5-E10.5 after cardioplegia in cold 250mM KCl (E9.5-E10.5), fixation in 4% PFA and dehydratation in methanol 100%. *Nppa, Pitx2, Wnt11, Bmp2, Fgf8, Tbx5, Cyp26c1, Hoxb1* riboprobes were transcribed from plasmids. Hybridization signals were detected by alkaline phosphatase (AP)-conjugated anti-DIG antibodies (1/2500), which were revealed with NBT/BCIP (magenta) substrate. After staining, the samples were washed in PBS and post-fixed. Brightfield images were acquired with a Zeiss AxioCamICc5 Camera and a Zeiss StereoDiscovery V20 stereomicroscope with a Plan Apo 1.0X objective.

Wholemount RNAscope ISH was performed with Mutliplex Fluorescent v2 Assay (Desgrange et al, 2020). E8.5 embryos were fixed 6h at room temperature or 24h at 4°C in 4% PFA, dehydratated in methanol 100%. *mm-Col5a1*, *mm-Greb1l*, mm-*Cacna1h, mm-Hapln1, mm-Fabp3, LacZ,* mm-*Nr2f2, mm-Aldh1a2, RNA 45S* probes were used, together with Hoechst as a nuclear counterstain. Amplification steps were performed using the TSA cyanine5 or 3 amplification kit. Samples were then transferred in R2 CUBIC clearing reagents. Multi-channel 16-bit images were acquired with a Z.1 lightsheet microscope and a 20X/1.0 objective.

#### Immunofluorescence

Immunofluorescence on whole mount E9.5 embryos was performed using CUBIC clearing as described in (Le Garrec et al., 2017) with Hoechst as a nuclear counterstain. Samples were blocked with the TSA Blocking Reagent (Perkin Elmer) with 0.5% Triton, incubated 48h at 4°C with the primary antibodies Acta2 (1/200) and Isl1 (1/30) and during 48h at 4°C with the secondary antibodies anti-mouse 647 (1/200) and anti-rabbit 488 (1/400). Multi-channel 16-bit images were acquired with a Z.1 lightsheet microscope (and a 20X/1.0 objective.

#### HREM (High-Resolution Episcopic Microscopy)

Embryos were collected, and after cardioplegia, fixed in 4% PFA. They were embedded in a methacrylate resin (JB4) containing eosin and acridine orange as contrast agents (Desgrange et al., 2020). One or two channel images of the surface of the resin block were acquired, using the optical high-resolution episcopic microscope and a 1X Apo objective, repeatedly after removal of 1.5-1.8 µm (E9.5) and 1.8-2.7 µm (E10.5, E11.5, E12.5, E13.5) thick sections: the tissue architecture was imaged with a GFP filter and the staining of enzymatic precipitates with an RFP filter. The dataset comprises 500-3500 images of 0.96-4.7 µm resolution in x and y depending on the stage. Icy and Fiji softwares were used to crop or scale the datasets. 3D reconstructions were performed with the Imaris Software.

#### RNA isolation

Collected embryos were staged based on the morphology of the heart tube(Le Garrec et al., 2017). The yolk sac was used for genotyping and sex determination (McFarlane et al., 2013) by PCR. Four embryos at E8.5e with the highest RNA quantity (43-159ng) were selected per genotype (*Greb1l^-/-^* mutants and *Greb1l^+/+^* wild-type littermates, *Greb1l^ex17/ex17^* mutants and *Greb1l^+/+^* wild-type littermates), with an equivalent sex proportion between controls and mutants of the same line. Micro-dissection was performed to isolate the cardiac region, from below the headfolds to the end of the second somite, after removal of the back. The tissue was flash frozen in liquid nitrogen. Total RNA was extracted in TRIzol-Chloroform and purified using the RNeasy micro kit including DNAse treatment. RNA quality and quantity were assessed by capillary electrophoresis using High Sensitivity RNA reagents with the Fragment Analyzer (Agilent Technologies) and RNA concentration was measured both by spectrophotometry using Fragment Analyzer capillary electrophoresis. All RINs were at 10.

#### RNA sequencing

RNAseq libraries were prepared starting from 50 ng of total RNA using the Universal Plus mRNA-Seq kit as recommended by the manufacturer. Briefly, the mRNAs were captured with polyA+ magnetic beads and fragmented chemically. Single strand and second strand cDNA were produced and ligated to Illumina compatible adapters with Unique Dual Index. The cDNA produced were PCR amplified after an initial test to evaluate the suitable number of PCR cycle to apply to each sample. To produce oriented RNAseq libraries, a final step of strand selection was performed. An equimolar pool of the final indexed RNA-Seq libraries was prepared, using the NuQuant system, and sequenced on a NovaSeq6000 from Illumina with paired-end reads of 100 bases and a mean sequencing depth of 50 millions per sample. Quality check of samples, for the genotype and microdissection is shown in Fig. S3A-C. The RNA-seq data are available in the NCBI Gene Expression Omnibus (GEO) database with the accession number (ongoing submission).

#### Phenotyping of congenital heart defects

Heart of *Greb1l* mouse mutants was phenotyped at E13.5, before in utero lethality, on 3D reconstruction of HREM images, based on the segmental approach (Van Praagh, 1972) and IPCCC ICD-11 Code.

#### Tamoxifen injection

*Tg(RARE/Hspa1b-cre/ERT2)#Asc R26^mTmG/mTmG^* males were crossed with B6D2 females. Tamoxifen was diluted in corn oil after 30 minutes heat (20mg/ml) and administered orally using feeding needles (Popper; cat No. 9921, size: 20GX1.1/2) at a concentration of 1mg per pregnant mouse, 6.5 days after the plug. Embryos were collected at E8.5 and cleared with a CUBIC approach(Le Garrec et al., 2017). Multi-channel 16-bit images were acquired with a Z.1 lightsheet microscope and a 20X/1.0 objective.

### QUANTIFICATION AND STATISTICAL ANALYSIS

#### Quantification of RNAscope ISH signal

The cardiac region was segmented in 3D images using the Imaris software. The heart tube and heart field were manually outlined at regular intervals of the Hoechst channel and the Create Surface function was used to reconstruct the 3D surface. The heart field was taken as the lateral plate mesoderm between headfolds and second somite, in agreement with fate maps (Domínguez et al., 2012). We used the Spot Detector tool to count the number of RNA molecules and calculate the heart tube/heart field ratio.

#### Quantification of the geometry of the heart tube at E9.5 and E10.5

From the 3D reconstruction of the myocardium contour, the axis of the cardiac tube was extracted, using eight landmarks along the length of the tube. Three of these landmarks were obtained with the Imaris Oblique Slicer function intersecting the tube perpendicularly, and by computation of the centroid of the polygon (Matlab geom3d library: function polygonCentroid3d) : one at the exit of the outflow tract, one at the sulcus between the two ventricles, and one at the bifurcation of the two atria. The five other landmarks were obtained by sub-division of the volume into the outflow tract, the right ventricle, the left ventricle, the atrioventricular canal and left atrium, and the right atrium, according to ISH labelling (E9.5) and anatomical landmarks such as cushion boundaries. The centroid of each of these volumes was computed, using the Imaris Software.

The 3D coordinates of the eight landmarks were used to do the quantifications. All the hearts were aligned. Two landmarks on the notochord were taken, as well as two dorsal-ventral landmarks in a single transversal plane, the neural groove and the most ventral point of the embryo, which were then readjusted at 90° with the notochord. Then, two successive 3D rotations were applied to the image coordinates, using an in-house Matlab code: first a rotation aligning the notochord with the Z axis, then a rotation aligning the dorsal-ventral axis with the X axis. All measurements were done on aligned samples. The origin for the 3D new reference axis was the distal outflow tract (exit of the tube) at E9.5 or a point localized at the bifurcation of the lung lobes at E10.5. The arterial pole was defined as the distal outflow tract and the venous pole as the bifurcation of the atria. The orientation of the RV/LV axis relative to the notochord, the distance between the poles along the antero-posterior axis and the tube length, were calculated as in(Le Garrec et al., 2017). The distance between the poles along the left right axis was calculated by subtracting the Y coordinates of the venous pole to the Y coordinates of the arterial pole in the reference axis system after alignment. The outflow tract length was measured by computing a distance between the centroid of polygons drawn along the outflow tract.

#### Quantification of the heart field size at E9.5

The heart field was segmented in 3D images using the Imaris software. The Isl1-positive heart field in the dorsal pericardial wall was outlined manually at regular intervals and the Create Surface function was used to reconstruct the 3D surface. The heart field was taken between the inferior wall the outflow tract and the junction with the atria. The size of the heart field is measured as the length along the cranio-caudal axis.

#### Quantification of criss-cross and supero-inferior ventricles

Analysis was carried out similarly on 3D images of E13.5 mouse and patient hearts, using Imaris and Matlab softwares. Planes of the interatrial and interventricular septa were identified anatomically and placed in 3D using an Oblique slicer. Similarly planes of the left and right atrioventricular valves were identified anatomically in 2 different views, and placed in 3D using an oblique slicer. Three points were annotated on the surface of each plane to define two vectors per plane. Then, each plane was defined by the normalized cross product of the two vectors, i.e. the orientation of the vector perpendicular to the plane. The angle between two planes was calculated as the arc-cosine of the dot product of the two vectors defining each plane.

Analysis of ventricle position is based on the orientation of the interventricular septum. The reference antero-posterior axis was defined by the spine in human and the notochord in mice. The reference dorso-ventral axis was taken in a single transversal plane, with one point on the notochord and one point on the most ventral point of the heart tube. It was then readjusted at 90° with the notochord using an in-house Matlab code. Both axes were defined by two points defining a vector. This vector was normalized. The reference left right axis was obtained by the cross product of the antero-posterior and dorso-ventral axis. Each 3D coordinates of ventricle position, reflecting the direction of the RV/LV axis, was obtained by the dot product of the normalized interventricular septum vector with each normalized reference axis vector (antero-posterior, dorso-ventral and left-right).

#### Computation of the torsion index

Following the hypothesis that the orientation of the ventricles may result from a rotation of the whole tube, we estimated the tube region most prone to it, as a consequence of a pulling force in the outflow tract. Considering the atria as fixed, if the various parts of the tube are approximated as cylindrical beams, the torsion angle resulting from the pull at the arterial pole would be 1) proportional to L, the distance from the centre of the region to the proximal OFT (where the pulling force is supposed to be applied), 2) inversely proportional to the quadratic moment of inertia of the beam I_G_=D^4^-d^4^, where D and d are the outside and inside diameters. The distance L and area of sections along the E10.5 tube were measured using the Imaris Software. The local torsion index was calculated as L/I_G_, providing a comparison of the amount of rotation likely to occur in distinct regions of the tube as a consequence of a pulling force.

#### Bioinformatics analyses of bulk RNA sequences

FASTQ files were mapped to the ENSEMBL Mouse GRCm38/mm10 reference using Hisat2 and counted by featureCounts from the Subread R package. Read counts, including duplicates (79,9% in mean), were normalized, and group comparisons were performed using the DESeq2 R package v1.24.0 (Love et al., 2014). Flags were computed from counts normalized to the mean coverage. All normalized counts < 20 were considered as background (flag 0) and ≥ 20 as signal (flag = 1). P50 lists used for the statistical analysis gather genes showing flag=1 for at least half of the samples. The model was adjusted for the effect of the condition (mutant or control, *Greb1l^-/-^* and *Greb1l^ex17/ex17^* lines) and for the effect of sample sex. Clustering of samples by genotype was controlled by hierarchical clustering using the Spearman correlation similarity measure and Ward linkage algorithm. For genotyping validation, reads of *Greb1l* exons 3 or 17 were aligned on IGV (Fig. S3C). The list of differentially expressed genes between controls and mutants was filtered at p-value ≤ 0.05. For the analysis of gene expression, normalized read counts below 250 were considered as background, based on expected low positive expression (*Fgf8*) and expected negative expression (*Neurod1, Myf5*). Genes were grouped in three categories depending on their p-value and corrected p-value (Benjamini-Hochberg procedure) for the MA plots. Functional analyses were carried out using Gene Set Enrichment Analysis (GSEA), with the CAMERA function from the limma R package v3.40.6 (Wu and Smyth, 2012) based on the same model as in the differential analysis. The interrogated gene signatures were from Gene Ontology Resource terms (GO terms) obtained from the Molecular Signature Data Base (Liberzon et al., 2015). The enrichment score is a two-sided t-test statistic as described in the CAMERA function from the limma R package v3.40.6 (Wu and Smyth, 2012). It measures how the genes in the set are highly ranked in terms of differential expression relative to genes not in the set. Heatmaps in Fig S3I-J show normalised gene expression centered and scaled by genes.

#### Bioinformatics analyses of single cell RNA sequences

Data from Tyser et al., 2021 was downloaded from https://marionilab.cruk.cam.ac.uk/heartAtlas/. The statistical analysis was reduced to cells from clusters Me3 to Me7, resulting in 1868 cells. Co-expression analysis to find genes associated with *Greb1l* was performed in two steps. The first step, adapted from de Soysa et al., 2019 consisted in applying the findMarkers function from scran R package v1.16.0 (Lun et al., 2016) to find genes differentially expressed between cells expressing (normalized counts > 0) and not (normalized counts = 0) *Greb1l*. The non-parametric test of Mann Whitney Wilcoxon was used, with the development stage added as a blocking factor, and both directions of expression were allowed (direction = ‘any’). The second step consisted in finding genes with normalized counts significantly correlated to *Greb1l* expression variations. Each candidate gene of the dataset was selected for correlation analysis whether it was expressed together with *Greb1l* in at least 100 cells: the pairwise Spearman correlation coefficient between normalized read counts was computed and its significance tested using an asymptotic t approximation. Corresponding p-values were adjusted for multiple testing using Benjamini-Hochberg correction (Benjamini and Hochberg, 1995). Cell cycle inference was performed using the CellCycleScoring function from Seurat package with the signature of cell stages provided in Seurat.

#### Generalized Linear Model (GLM)

In order to assess whether some geometrical parameters of the heart tube may explain the orientation of the RV/LV axis, a circular GLM model was used. This was implemented with the R package CircGLMBayes (Mulder and Klugkist, 2017), and the Shiny graphical user interface. The Shiny analysis options were: Iterations (10000), Burnin (1000), Link function range (0.51 as the RV/LV axis orientation amplitude is 90°). The outcome variable was the RV/LV axis angle, and various continuous or categorical predictors were tested (continuous: outflow tract length, left-right distance between the poles, antero-posterior distance between the poles; categorical: genotype), together with combinations of these predictors. The model fit was assessed using the Akaike Information Criterium (AIC), output from the Shiny analysis, and the AICs of the different models were compared. The relative likelihood of the various models compared to the best model was computed as exp((AIC_b_ – AIC_i_)/2), where AIC_i_ is the AIC of the assessed model and AIC_b_ is the AIC of the best model.

#### Statistical analysis

Collection of full litters was used to randomise imaging experiments. Group allocation was based on PCR genotyping. All sample numbers (n) indicated in the text refer to biological replicates, i.e. different embryos or different cells. Investigators were blinded to allocation during imaging and phenotypic analysis, but not during quantifications. Tests were performed with GraphPad Prism and R. Correlation between two data series was quantified by the square of the Pearson coefficient R^2^. Comparisons of two central tendencies were done on the mean using a Student two-tailed test. Angle comparison of the RV/LV axis, valve planes and ventricle position were compared using a non-parametric Mann-Whitney test. When more than two central tendencies were compared, an ANOVA was calculated with Tukey multi-comparison tests. One sample control at E9.5 and one sample at E10.5 were considered as outliers in the geometric analysis and excluded because of a growth delay based on the somite number and the length of the heart tube.

### RESOURCE AVAILABILITY

#### Lead contact

Further information and requests for resources and reagents should be directed to and will be fulfilled by the Lead Contact, Sigolène M. Meilhac

#### Material availability

This study did not generate new unique reagents.

All stable reagents generated in this study are available from commercial sources or the Lead Contact without restriction

#### Data and Code availability

The RNAseq dataset generated during this study is available at the NCBI Gene Expression Omnibus (GEO) database with the accession number (ongoing submission).

